# Auditory brainstem responses to speech-in-noise reflect selective attention, comprehension, and subjective listening effort

**DOI:** 10.1101/2024.12.23.629710

**Authors:** K. Mankel, D. C. Comstock, B. M. Bormann, S. Das, D. Sagiv, H. Brodie, L. M. Miller

**Author notes:** **Corresponding author:** Kelsey Mankel, 4055 N Park Loop, Memphis, TN, 38120.

## Abstract

The auditory brainstem plays a crucial role in speech-in-noise listening, refining numerous acoustic features such as pitch and spatial location under continual descending influence from cortex. However, the difficulty of characterizing brainstem activity during continuous speech listening has obscured its functional role in ecologically valid contexts—not only the effects of selective attention on neural responses, but also their impact on comprehension and listening effort. Here, we evaluate the role of the brainstem on speech-in-noise perception and selective attention using a continuous, speech-based stimulus with embedded chirps (Cheech) that rapidly and effectively evokes auditory brainstem responses (ABRs) while participants listen to short story narratives. The Cheech-modified stories were presented alone or in the presence of another spatially separated talker, and neural responses were measured throughout by EEG. Both word-level detection performance and narrative-level comprehension were evaluated, as well as subjective reports of listening effort. ABR wave V was modulated by the presence of a competing talker, selective attention for the target versus masker, and the talker sex. Additionally, faster wave V peak latencies and larger amplitudes were associated with identification of the target words embedded within the stories, while faster latencies additionally related to better comprehension question accuracy and lower subjective listening effort. Collectively, our results provide evidence for the influence of brainstem encoding processes on individual speech-listening behaviors, including the ability to selectively attend to and comprehend target speech in the presence of a competing talker.

**Significance statement:** This study highlights the crucial role of the brainstem in speech-in-noise perception and higher-level language abilities like comprehension and subjective listening effort. Through the use of a novel speech stimulus blended with embedded chirps (Cheech) to evoke brainstem responses while listening to short story narratives, we show how selective attention influences subcortical processes. Faster and more robust subcortical processes, in turn, were associated with more successful speech-listening performance across multiple behavioral metrics. These findings advance our understanding of how the brainstem supports selective attention and speech perception in challenging listening environments.

## Introduction

Neural processing of speech has primarily been studied at the level of the cortex (e.g., Hickok and Poeppel, 2004). Yet, brainstem nuclei along the auditory pathway perform crucial functions relevant for processing speech such as encoding pitch and timing of spectrotemporal fluctuations (e.g., Krishnan and Gandour, 2009; Krishnan and Plack, 2011), preliminary phoneme categorization (Carter and Bidelman, 2023; Rizzi and Bidelman, 2023), and even early differentiation of competing auditory streams, likely through corticofugal modulations (Price and Bidelman, 2021). Additionally, poorer encoding of auditory brainstem responses (ABRs) has been associated with decreased speech-in-noise (SiN) performance (Papakonstantinou et al., 2011; Bramhall et al., 2015). Thus, rather than serving as a simple relay for acoustic information, the brainstem plays an active, perceptually consequential role in speech perception.

Yet, whether and how subcortical processes influence the ability to selectively attend to target sounds while filtering out background noise, as required for SiN perception, remains uncertain. In early studies, ABRs were considered immune from effects of attention and arousal states (e.g., Jewett and Williston, 1971; Connolly et al., 1989; Gregory et al., 1989). However, midbrain responses (ABR wave V) may be modulated by selective attention of audition over another modality (Lukas, 1980; Bauer and Bayles, 1990; Kumar et al., 2023), increased cognitive load (Sörqvist et al., 2012), or cognitive interference (Brännström et al., 2020). Mixed results on the influence of attention have also been demonstrated for other subcortical responses like the frequency-following response (FFR) (e.g., Hoormann et al., 2000; Galbraith et al., 2003; Varghese et al., 2015), possibly due to cortical contributions (Coffey et al., 2016; cf. Bidelman, 2018). Meanwhile, regression- or correlation-based brainstem responses derived from continuous speech are more robust for attended rather than unattended talkers (Forte et al., 2017; Etard et al., 2019), and attentional modulation of these responses relates to independent tests of SiN perception (Saiz-Alía and Reichenbach, 2020).

Historically conflicting results regarding attentional modulation of brainstem activity likely reflect differences among experimental approaches, chief among them a varying degree of ecological validity. While subcortical EEG responses are best evoked by transient artificial stimuli such as isolated clicks or tone bursts, these are not always good predictors of speech perception in real-world listening environments (Skoe and Kraus, 2010), nor do simple sounds adhere to naturalistic statistics known to influence midbrain encoding (Escabí et al., 2003). As a result, despite numerous studies of auditory brainstem function, its role in attentive listening and its relevance to real-world speech performance remain unclear. In order to shed light on the neural processes underlying SiN perception, a challenging, ecologically valid task using naturalistic speech with relevant outcome metrics should be used (Hamilton and Huth, 2020).

However, fundamental limitations characterize ABR recordings such as low signal-to-noise ratios (SNR), relatively small amplitudes, and the need for multiple stimulus repetitions. To address this, we have developed a technique for the rapid measurement of brainstem through cortical responses via “chirped-speech” (Cheech), continuous speech stimuli blended energetically and perceptually with frequency sweeps (Backer et al., 2019; Miller et al., 2020; Shehabi et al., 2025). Our technique parallels recent efforts to derive brainstem activity during continuous speech (Forte et al., 2017; Maddox and Lee, 2018; e.g., Etard et al., 2019; Polonenko and Maddox, 2021; Kulasingham et al., 2024). Compared to other methods, Cheech offers more acoustic control over the specific parameters used to elicit multiple, simultaneously recorded brain responses from naturalistic speech, all while participants perform a challenging listening task complete with spatial cues and attentional switching.

Here, we use Cheech-modified short stories presented both in quiet and in the presence of a competing talker to elucidate attentional effects within the brainstem through ABRs (i.e., wave V). Our results suggest wave V is affected not only by the presence of another talker, but also by selective attention to one talker over another. We also demonstrate the relationship between ABR wave V and several higher-level speech listening behaviors, including comprehension and subjective listening effort.

## Methods

### Participants

Twenty-nine participants were recruited for this study. To be eligible, prospective participants must be between 20 and 40 years old, speak English as their first language, and report no neurologic, psychiatric, or scalp conditions that directly affected their ability to understand speech or interfere with electrophysiologic measurements (e.g., epilepsy, certain strokes, ADHD, scalp problems, etc.). Participants were screened for visual acuity (i.e., better than 20/40 vision per Snellen chart), normal hearing (i.e., ≤25 dB HL air conduction thresholds between 250-8000 Hz in both ears, no air-bone gaps >10 dB within 500-2000 Hz, no asymmetries between the ears >20 dB at 500, 1000, or 2000 Hz), and cognitive abilities (>24 Montreal Cognitive Assessment [MoCA] score—consistent with an ongoing comparison study group of veterans; Nasreddine et al., 2005). One participant withdrew from the study due to personal reasons; one participant was excluded due to technical difficulties during the EEG recordings, and two participants were excluded because they did not pass the hearing screener at the time of testing. This resulted in a final cohort of N=25 total participants with the following demographics: average age=22.88 ± 4.98 SD years old (range=18-38 years old); 15 females and 10 males; N=21 self-identified handedness right, N=2 left, and N=2 ambidextrous. Participants had on average 16.86 ± 3.19 SD years of education and 4.20 ± 3.74 SD years of formal music training. Of those who reported some level of formal music training (N=22), the average age they began music training was 10.05 ± 2.75 SD years old. Average MoCA score was 28.16 ± 1.46 SD (range=25-30). Additionally, participants completed a version of the MacArthur Scale of Subjective Socioeconomic Status where they were asked to rate their perceived socioeconomic status (SES) relative to individuals across the US society using a ladder-rung rating from 1 (lowest SES)-10 (highest SES) (Adler et al., 2000). Mean subjective SES scores were 4.70 ± 1.30 SD, range 3-8. Average air conduction pure tone average (PTA) thresholds were 6.80 ± 3.72 SD and 7.04 ± 4.47 SD for the right and left ears, respectively, indicative of normal hearing. All participants completed an informed consent as approved by the University of California, Davis Institutional Review Board and were compensated financially for their participation.

### Stimuli

The stimuli were short fairytale stories available in the public domain (e.g., on www.librivox.com). The stories were selected for content unlikely to be widely known, to avoid comprehension question recall based on modern renditions (i.e., popularized by children’s books or movies), and with story arc progress within a 7.5-minute block (i.e., rising action, climax, or falling action) to ensure sufficient and generally consistent content for probing comprehension. Thirteen stories were piloted on four raters who scored them according to factors such as interest and engagement levels, ease of understanding, grammar or writing style complexity, familiarity (i.e., low scores preferred), and suitability for an adult audience. Seven stories that scored highest across these factors were ultimately retained for the experiment. For more information about the specific stories used in the current study, see Extended Data Table 1-1.

The short stories were modified to add monosyllable color words as targets for the spatial attention experiment (e.g., red, green, blue, white, etc.). The color words were embedded as part of the story itself rather than unassociated, random target word interjections (e.g., “…with a narrow ***<u>blue</u>***ribbon over her shoulders”). We recorded a male talker (native English speaker with a neutral American accent) reading the modified short stories aloud using a Shure KSM244 vocal microphone (cardioid polar pattern, high-pass filter of 18 dB per octave cutoff at 80 Hz) and Adobe Audition (sample rate = 48,000 Hz, 32 bit depth [float]) in a quiet, soundproof room. Silent gaps in the recordings greater than 500 ms were shortened to 500 ms, similar to (Broderick et al., 2018; Teoh and Lalor, 2019; Teoh et al., 2022), and then the trimmed recordings were then cropped to 7.5 minutes per story audio.

To produce dual-talker conditions, one story from each pair was pitch modulated using MATLAB STRAIGHT prior to the Cheech process (see Chirped speech (Cheech) below). This manipulation allowed us to compare neural and behavioral responses for male versus female voices (between-story comparison) in addition to attended versus unattended conditions (within-story comparison; see Experimental conditions section). Specifically, female-intended stories were modulated in pitch to an average F0 of 180 Hz and modulated in frequency to model a shorter vocal tract length, while the original (male voice) audio had an average F0 of approximately 128 Hz. Modifying voice parameters in this manner meant speaker characteristics such as phrasing, intonation, and pacing were otherwise preserved because all stories were originally spoken by a single talker. Modifying one talker’s F0 also provided the added benefit of a combined voice-sex + spatial release from masking which improved pilot test performance (Oh et al., 2021, 2022).

### Chirped speech (Cheech)

The short story auditory stimuli were then modified via a patented process called “Cheech” (“chirped-speech”) to evoke the auditory evoked potentials (Backer et al., 2019; Miller et al., 2020; Shehabi et al., 2025; Figure 1). In Cheech, some of the glottal pulse energy within predefined frequency bands are replaced with synthetic chirps, consistent with previously published protocols from our lab (Backer et al., 2019). Aligning the chirps with glottal pulse energy inherent in speech creates an acoustically-fused auditory perceptual object with sufficient transient chirp activation to measure auditory brainstem through cortical responses while preserving naturalistic speech qualities and linguistic content. First, periods of sufficient voiced power were identified; specifically, the audio was filtered from 20 to 1000 Hz, and voiced periods at least 50 ms long were identified when the speech envelope between 20-40 Hz surpassed a threshold ∼28% of overall speech root-mean-square (RMS) amplitudes. The timing of the glottal pulses during voiced periods was determined by a speech resynthesis process using custom MATLAB code and the TANDEM-STRAIGHT toolbox that retains original speech frequency characteristics (Kawahara et al., 1999, 2008; Kawahara and Morise, 2011). The continuous speech was re-filtered into alternating, octave-wide frequency bands of 0-250, 500-1000, 2000-4000, 11,000-∞ Hz, and chirp energy was constrained to frequency bands of 250-500, 1000-2000, 4000-11,000 Hz. The chirp and speech bands in the alternating, interleaved octave-wide bands were then added together. Each mono track was then duplicated to form stereo audio. Audio files were normalized prior to the Cheech synthesis to ensure equivalent chirp amplitudes relative to the speech levels across each story, and the root-mean-square (RMS) amplitudes were re-checked across all Cheech-ed stories for equivalence after the Cheech process was completed.

**Figure 1:**
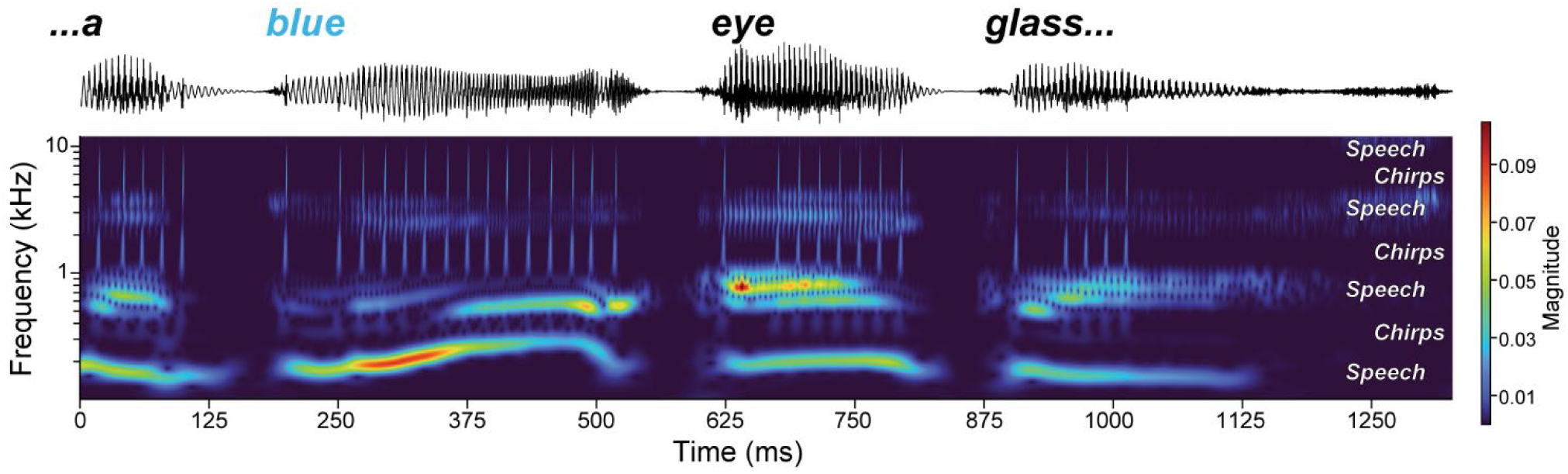
A brief segment of the chirped-speech (Cheech) stimuli. (Top) Participants listened to the Cheech-modified short stories and pressed a button when they heard the target talker speak a color word (e.g., “blue”). (Middle) Stimulus waveform and (bottom) spectrogram for the Cheech audio. Cheech interleaves speech and chirp frequency bands in a manner that preserves naturalistic speech qualities while evoking robust ABRs. See Methods for details.

Timing of the chirps varied with the natural fluctuations in the running speech, with special modifications to optimize measurement of the auditory ERPs. Specifically, when voicing energy was detected, chirps were inserted at minimum 18.2 ms apart (55 Hz), skipping glottal pulses that occur within the 18.2 ms window. The first chirp within a sequence is always followed by a minimum of 48 ms before the next chirp is presented, skipping glottal pulses in between, to produce a longer ISI that allows for improved evolution of middle latency responses (MLR). For a summary of specific Cheech processing details, see Extended Data Table 1-2.

### Spatial filtering and attention switching task

One goal of this study was to design a paradigm that mimicked real-life communication, including the ability to switch spatial attention to different talker locations. Spatial attention has been shown to modify neural activity associated with stream segregation (e.g., alpha power band lateralization), but these effects are often observed hundreds of milliseconds later than the ABRs explored here (e.g., Kerlin et al., 2010; van Diepen et al., 2016; Wöstmann et al., 2016; Teoh and Lalor, 2019). The results associated with spatial attention differences will be presented in a later publication. For completeness, we describe the details of the spatial attention switching task below, but for the sake of current analyses, results were collapsed across the attend-left and attend-right conditions.

The Cheech stimuli were spatially filtered via HRTFs to simulate audio at ±15° (SADIE II database; Armstrong et al., 2018). Spatial audio was mirrored by two symbols on two computer screens approximately 15° to the left and right of the midline in front of the participant. A “+” symbol on one of the screens corresponded to a given target location. “<” and “>” symbols were shown on the computer screen opposite from the target speech location (e.g., a “>” shown on the left screen and a “+”on the right screen indicated a the target voice located in the right auditory spatial field; see Figure 2). Consequently, “<” and “>” symbols indicated either the absence of another speaker for a given spatial field, as in the mono-talker condition, or the location of the masker story in the two-talker condition. The symbols were identical in size, color, and line lengths to maintain equivalent luminance for the eye tracking measurements (data not included here). The spatially filtered audio and visual symbols were then edited with custom MATLAB code to swap left and right spatial locations 75 times, or approximately every six seconds (7.5 minutes ÷ 75 switches = 6 seconds; switches assigned per random distribution of 6 ± 1 seconds. The visual icons switch screens instantaneously at each switch time while the crossover fade time for the audio was 35 ms, beginning at the onset of a switch cue (Figure 2). During piloting, this crossover time was an appropriate balance between minimizing “pops” and other artifacts from cutting the audio across left and right audio tracks while also minimizing the perception of a gradually moving sound between spatial locations (i.e., a sudden jump from one side to the other without introducing robust distortions). In this manner, our spatial attention switching task mimicked listening scenarios with multiple alternating targets, as if perceptually following a conversation between two spatially separated talkers.

**Figure 2:**
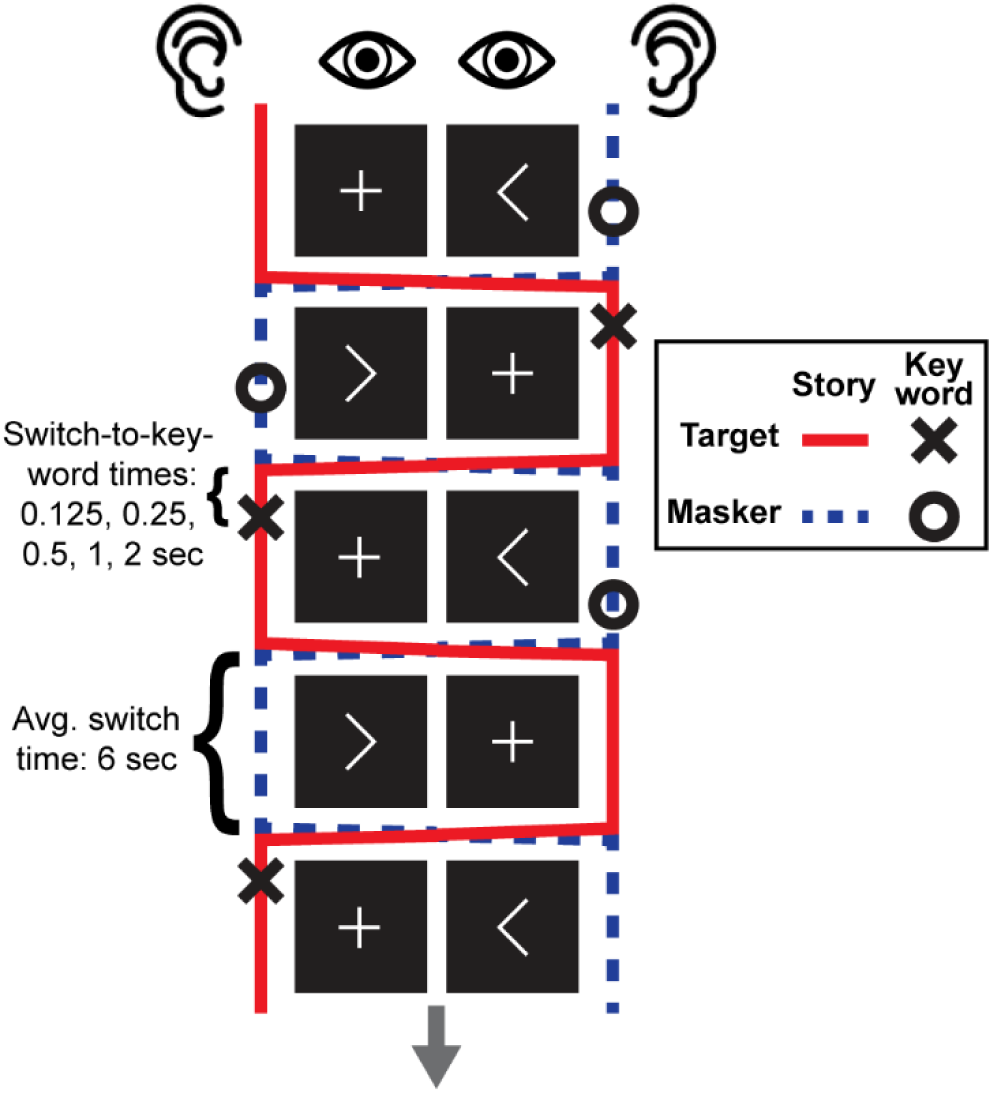
Experimental paradigm. Participants are instructed to attend to the target speech (red line) and press a button when a color word is spoken (‘X’). They must ignore color words spoken by the masker talker when present (i.e., dual talker condition; ‘O’). The target visual cue and voice alternate between approximately +/- 15 ° in the visual and auditory spatial fields on average around every 6 seconds (see Methods).

Color words were randomly assigned to one of five possible onset categories between 0.125 and 2 seconds from the switch-to-color-word-onset (i.e., 0.125, 0.25, 0.5, 1, and 2 seconds, 5 color words per condition; see Figure 2). These categories were established after preliminary piloting to assess whether “attentional blinks” from switching left to right spatial attention following the cue (and vice versa) impaired color word detection across participants; these data will be presented in a subsequent paper. Onset timings of the color words were determined by Gentle, an audio forced aligner (https://github.com/lowerquality/gentle). The closest preceding switch time was replaced by the assigned switch-to-color-word-onset, resulting in 50 random switches and 25 switch-to-color-words (i.e., 33% probability of a target color word onset within 2 seconds of a switch). The 25 words were balanced across left-to-right and right-to-left switches within a story (minus one remainder).

### Experimental conditions

Following audio cropping to 7.5-minute length, the seven stories were then divided into separate experimental conditions: two stories presented alone (“mono-talker”), two sets of paired stories (“dual-talker”), and one for initial practice which was not included in subsequent analyses. Within the dual-talker condition, each story was heard twice—once as the “target story” and once as the “masker story” within the pair. This experimental manipulation thus allowed us to measure effects of directed spatial attention because the same audio was presented and participants were instructed to either “attend” or “ignore” the story, respectively. Mono-talker stories were only presented once each. The target story always started on the left side, whether presented alone (mono-talker) or in the presence of a masker (dual-talker). The order of stories and conditions was counterbalanced with a Latin square design across participants to reduce potential sequence-dependent learning effects. No story pair was heard back-to-back to reduce short-term learning effects as well.

Story pairs in the dual-talker condition were subject to additional considerations. As mentioned in the Cheech section, one voice within the story pair was pitch shifted to simulate a female speaker. Consistent with combined voice-sex and spatial release from masking outcomes observed in previous work (Oh et al., 2021, 2022), two distinct talker F0s helped listeners better distinguish between the spatially-separated voices and more successfully complete the task during piloting. The mono-talker stories were presented in the original, unmodified male voice only. Initially, female mono-talker stories were planned as well, but it became clear during piloting that additional stories would significantly increase the total testing time and lead to excessive fatigue, so these stories were ultimately dropped from the final experimental design.

Given the constraint of naturalistic color word embeddings within each story, color words occasionally overlapped within the dual-talker pairings. The story pairs were determined according to those which had the fewest color word overlaps, and any overlapping color words were dropped from the behavioral task analyses so behavioral responses to target (hits) and masker color words (false alarms) would not be conflated. A few target story switch times were also manually adjusted to ensure that no masker color word occurred during a switch (i.e., masker color word onset was not between -0.5-0.1 ms from a [target story-driven] switch time).

Additionally, half of the stories—one mono-talker story and a pair of dual-talker stories— were subjected to additional modifications during the Cheech process. These modifications were specifically intended to disrupt subcortical encoding mechanisms, so we do not include analysis of those stories here. These data will be analyzed in separate reports. Rather, this study focuses on the remaining 3 experimental stories (i.e., one mono-talker story and a pair of dual-talker stories). For additional details on the short stories used in the current study, see Extended Data Table 1-1.

### Stimulus presentation equipment

Participants were seated in a soundproof, electromagnetically shielded booth (ETS Lindgren), approximately 176 cm from two computer monitors. The Dell UltraSharp monitors had a refresh rate of 60 Hz, and screen settings were left on standard factory defaults. Stimulus presentation, including both audio and visual, was controlled by custom MATLAB code using Psychtoolbox (Brainard, 1997; Pelli, 1997). Auditory stimuli were routed from a Hammerfall audio card to an audio interface (RME Fireface UFX II), then to a headphone driver (RME ADI-2DAC FS) which drove ER-2 insert earphones (Etymotic Research/Interacoustics). The ER-2 inserts were electromagnetically shielded to reduce EEG artifacts (Campbell et al., 2012). Cheech stimuli were presented at 75 dB SPL because conversational speech levels are often moderately elevated in the context of noisy settings (Rusnock and Bush, 2012). Clicks for the separate ABR recordings were presented at 80 dB SPL. Stimuli were calibrated using a Larson-Davis PRM902 preamp & 2221 power supply, B&K 4157 coupler, and C-weighted average.

### Procedure

In total, each participant completed three sessions as part of a larger experiment: one 1-hour audiologic exam, one 2-hour behavioral session, and one 2.5-hour laboratory EEG session. Several participants completed more than one session on the same day (i.e., either audiologic + behavioral or behavioral + EEG sessions), but a long break was offered in between sessions to reduce fatigue. The data presented here focuses on the EEG experimental session only.

The order of events within each block is as follows: 1 minute of brief rest (for resting state EEG analysis), 7.5 minutes of the spatial attention listening task, 5 comprehension questions, 6 subjective task load questions (Hart and Staveland, 1988), a flash reaction time task (5 short trials), and then a break. During the one-minute rest period, participants were instructed to remain still with their eyes open. During the spatial attention listening task, participants listened to the short stories, following the target story and corresponding visual cue as they switched between left and right relative spatial locations (Figure 2). They were instructed to press the spacebar when they heard the embedded color words within the story. In dual-talker condition blocks, participants were told to ignore color words from the masker story and only press the spacebar for target story color words. The audio was immediately followed by a series of five multiple choice comprehension questions with four possible answer choice options (chance = 25% accuracy). Comprehension questions were balanced for difficulty (not too hard nor too easy; piloted for accuracy on 3-5 individuals per story), story time relevant for the question (i.e., early vs. late in the 7.5-minute story segment), and memorization vs. inductive reasoning probed by the answers. Next, participants completed the NASA Task Load Index, a measure of subjective task-related workload or effort, modified for computerized data collection using a sliding scale selection tool (Hart and Staveland, 1988). To measure general mental fatigue across the blocks, we used a psychomotor vigilance-based visual reaction time task (e.g., Basner & Dinges, 2011; Dinges & Powell, 1985; Hornsby, 2013; Wilkinson & Houghton, 1982). Participants were instructed to watch the crosshairs shown at the center of the computer screen and press the spacebar as quickly as possible when a white circle appears on screen. The circle flashed on screen for 40 ms across five trials. Finally, participants were offered a break before starting the next block. The entire experiment presentation was controlled by custom MATLAB and Psychtoolbox code.

The data presented here were collected as part of a larger study. As mentioned in the Experimental Conditions section, participants completed three additional short story blocks with Cheech audio designed to impair auditory brainstem encoding, so those blocks are not included in the present analyses. Additionally, participants obtained a full audiologic evaluation and completed a battery of cognitive tests in separate experimental sessions. These data will be analyzed in separate reports. Many participants completed more than one session in a given day but not all three on a single day, and they were given longer breaks in between sessions to reduce fatigue.

### Behavioral outcome measures and analysis

Selective attention listening was assessed by correct identification of color words embedded in the target story (hits). A “hit” was defined as pressing a designated keyboard button within 2 seconds after the onset of a target word. Any pairs of color words within +/-2 second onset across the target and masker stories in the dual talker condition were excluded from analyses, resulting in four overlapped color words dropped from each story in the behavioral task analyses for the pair of stories used here. Specifically, the relevant measures were the proportion and corresponding reaction times of correct hits out of the total possible, non-overlapping target words (n=25 in the mono talker condition, n=21 per story in the dual talker condition). Behavioral outcomes measured after each story concluded were comprehension question accuracy (i.e., total % correct out of five questions), task load effort scores (i.e., average NASA task load index scores across six questions, where higher scores means greater perceived task load; Hart and Staveland, 1988), and average reaction times on the vigilance-based flash task in a given block.

### EEG recordings

EEG recordings were obtained using a BioSemi ActiveTwo system (BioSemi B.V., Netherlands, www.biosemi.com) in an electrically shielded booth with a sample rate of 8192 Hz recording data from 72 electrodes: 64 channels following the International 10-20 standard with 4 additional channel pairs around the ears (earlobe, mastoid, lower posterior: posterior to the earlobe and inferior to the mastoid, and upper anterior: superior-anterior to the tragus near the zygomatic arch). Electrode offsets from all channels were measured to be below 20 µV relative to the Common Mode Sense electrode using ActiveView2 software. Data were recorded using open-source Lab Streaming Layer (LSL) software (https://github.com/sccn/labstreaminglayer) as XDF files on a desktop computer running Windows 10.

A pair of channel outputs containing trigger events was routed from the audio interface to a pair of Brain Products StimTraks for monitoring triggers associated with the target and masker stories, respectively, which then convert the audio trigger signals to trigger pulses sent to the TriggerBox (Brain Products GmbH). This setup permitted audio playback synchronized with chirp triggers embedded during the Cheech process with minimal latency delays (see Cheech section); such near-simultaneous synchronicity is crucial for measuring rapid, lower-level auditory brainstem responses during the listening task. Triggers controlled by Psychtoolbox (i.e., participant responses, block sequence triggers, spatial location switches, etc.) were also routed to the TriggerBox via the computer parallel port. The TriggerBox then passed both triggers from the parallel port and the StimTraks to the Biosemi signal receiver box.

### EEG Preprocessing

EEG were preprocessed using MATLAB 2021b (MathWorks Inc.) using a combination of EEGLAB functions (Delorme and Makeig, 2004), ERPLAB functions (Lopez-Calderon and Luck, 2014), and custom MATLAB code. Data were first imported into EEGLAB with events extracted from the trigger channel in the XDF file using custom MATLAB code. Given the large number of tightly packed event codes for our experimental design (>130,000/participant at an average rate of 1 event every 20ms), event codes from the 3 separate streams (parallel port and 2 StimTraks, described above) occasionally arrived at the TriggerBox simultaneously, resulting in correctable event code errors. Custom code was used to check the recorded events against the expected events for a given condition to add missing events, remove extra events, and adjust event latencies for any overlapped event. A second set of corrections were then applied to adjust for latency differences between the acoustic signal and the event times. These corrections took into account a 1ms delay due to audio travel time from the insert earphone tubes, a 1.973ms delay to the audio signal due to the HRTF process, a 0.541ms delay to the chirp triggers to account for a difference between the expected chirp timing from the Cheech process and the chirp onsets, and a 1ms delay in one of the StimTrak channels caused by the TriggerBox hardware.

Data were then pruned to retain only rest periods and attentive listening task periods along with a 10 second buffer on both sides of each selected period. Following pruning, each section of the data was DC corrected and a 2^nd^ order 0.1 Hz high-pass Butterworth filter was applied to remove slow drifts. We then referenced the data to electrode Cz. Even though our data were collected in an electromagnetically shielded booth, we occasionally found 60 Hz line noise in some channels. To minimize the effects of cleaning line noise, only channels with detected line noise were cleaned (mean = 4.4 channels ± 7.56 SD, range for a given participant = 0-39 channels). This was carried out by first checking for line noise using a frequency tagging approach. Spectra were calculated across all frequencies for each channel. For each frequency, the average spectra of 4 neighboring frequencies except those immediately adjacent (x-2, x-3, x+2, x+3) was subtracted from the spectra for each frequency. If the neighbor corrected power at 60 Hz or any of its first four harmonics (120, 180, 240, 300 Hz) was greater than 8 standard deviations calculated from the corrected power from remaining frequencies, that channel was cleaned of line noise using the Cleanline plugin (https://www.nitrc.org/projects/cleanline). This process was repeated until no further line noise was detected up to a maximum of 5 repetitions.

To clean the data of prominent artifacts such as eye blinks, eye movements, and heartbeat signals, we used Independent Component Analysis (ICA) following the methodology outlined by Luck (2022) which improves ICA performance, and greatly reduces the required processing time, compared to more traditional approaches. This process involved making a copy of the processed data (the ICA dataset), down-sampling and pre-cleaning it of prominent (non-eye and heart) artifacts, applying ICA to select and remove eye and heart artifacts, then transferring the calculated component weights back to the original dataset. Specifically, we first down-sampled the data to 256 Hz to speed ICA processing, then we inspected and rejected bad channels by eye (mean=3.72 ± 2.13 SD channels, range=0-9 channels). Following channel rejection, we applied a non-causal 8^th^ order Butterworth band pass filter with cutoffs at 1 and 50 Hz. Further cleaning was then undertaken using ERPLABs continuous artifact rejection function to remove sections of data with significant movement and muscle amplitude artifacts, the presence of which can hinder ICA’s ability to isolate eye and heart components (mean amount of data in seconds removed for ICA=129.19 ± 86 s SD or 3.6% ± 2.35%, range=4.98-298.79 s or 0.14-7.96%). The rejection criterion operated on a 1-s sliding window with a 0.25 s step size and an initial threshold of 200 µV. As we wanted to retain eye activity for ICA, frontal channels that contain the largest eye related activations (FP1, FPz, FP2, AF7, AF3, AFz, AF4, AF8) were excluded from the rejection threshold. We then inspected and adjusted the threshold level for each dataset (mean=220 ± 28.87 SD µV, range=200-300 µV) until only those sections of data with large muscle or movement artifacts were rejected while any eye blink or eye movement related artifacts remained. To further speed the ICA processing time, the number of input channels was reduced from an average of 67 channels to the first 32 principle components using Principle Component Analysis (PCA), then ICA was performed using the infomax algorithm. Eye blink, eye movement, and (when present) heart artifact components were manually marked for rejection (mean=2.8 ± 0.65 SD components, range=2-4 components). Following ICA processing, we prepared the original dataset for transferring component weights by first removing the same bad channels marked for removal in the ICA dataset. We then transferred the calculated ICA weights from the ICA dataset to the original dataset and rejected the marked artifact components in the original dataset. Thus, any eye and heart artifact components were removed from the dataset with the original sample rate (and without the aggressive pre-cleaning done to prepare the ICA dataset).

Once the data was cleaned with ICA, we applied a non-causal 8^th^ order Butterworth bandpass filter with cutoffs at 100 and 1500Hz and then re-referenced to the average of the left and right earlobes. If one of the earlobe channels had been rejected as a bad channel, the average of the left and right mastoid was used instead. Activity at channel Cz (the previous reference) was then recalculated and added back to the dataset. We then interpolated any rejected bad channels using spherical interpolation.

Epochs were extracted between –2ms and +15ms of chirp onset for all conditions with baseline correction from –2 to 0ms. The number of epochs was balanced so that each condition had the same number of chirps both in total, and from each spatial location (left, right), resulting in each condition (mono talker male, dual target male, dual target female, dual masker male, dual masker female) having 7,886 epochs per condition equally distributed across left and right presentation (3,943 epochs from both left- and right-side presentations per condition). The smallest number of chirps on any one side for all conditions was calculated and epochs were randomly sampled so all conditions were balanced. Random sampling and balancing the number of trials reduced possible transient effects of attention-driven modulations during or immediately after a spatial attention switch. Additionally, balancing of the number of epochs for each condition was done to avoid any systematic condition differences due to a difference in signal to noise (SNR) caused by some conditions having more chirps than others. This difference was primarily due to the female-voiced stories, where more frequent glottal pulses (driven by a higher F0) resulted in approximately 9% more chirps in the female-voiced stories compared to the male-voiced stories. Additional differences in total chirp events were due to an approximate 3% variation of chirps across stories (i.e., subtle differences in voicing and pausing). Given evidence for left-right asymmetries in speech processing and neural encoding (e.g., “right-ear advantage”; Ahonniska et al., 1993; Jerger and Martin, 2004; Hornickel et al., 2008; Keefe et al., 2008; Bidelman and Bhagat, 2015), we also balanced the number of chirps across stimulus presentation side for each condition to ensure results were not driven by unequal ear-based SNR or ear presentation differences in the EEG measures (approximately 5% variation in the of number of chirps across sides within each story). Epochs with artifacts (e.g., muscle activity) were then removed with a 2-step process: first, any epochs with activity outside the threshold of +/-100µV were rejected, and then any remaining epochs whose activity was outside of 6 SD for a single channel or 2 SD across all channels were eliminated (mean=9.85 ± 6.24% SD of rejected epochs, range=3.65-29.96% rejected epochs across individuals and conditions).

ABR Wave V peak amplitudes and latencies were extracted at channel Cz from the average amplitude over time across the remaining epochs for each condition for each participant. Wave V latency measures were extracted by fitting a local positive peak in a window between 5-8 ms post chirp-onset. Wave V amplitude was computed as the peak-to-peak amplitude between local positive peak within 5-8 ms and the subsequent local negative peak between 7-11 ms. The measurement windows were determined based on visual inspection of all individual Wave Vs for each condition. In cases in which more than one local peak was seen in the wave V waveform inside the measurement window, the values from the peak with the greatest amplitude were taken.

### Statistical analyses

Unless otherwise specified, mixed effects linear regression models and repeated-measures ANOVAs were used to analyze the data in SAS Studio (v.3.81). All regression models included a random intercept of subject to account for inherent variability present between individuals and improve generalizability of the results. Outcome measures were ABR wave V amplitudes and latencies for the neural models, and the behavioral metrics (i.e., target word identification accuracy, target word identification reaction time, comprehension question accuracy, subjective task effort ratings) were included as dependent variables for the behavior-only models as well as the brain-behavior comparisons. Independent variables were characterized by the condition of interest (i.e., one versus two talkers; target versus masker; also main effects and interaction terms with male versus female voices where appropriate). When interactions were not significant in the full model, they were removed using a backwards selection procedure, and the remaining simplified model variables are reported in the results below. One-way repeated-measures ANOVAs were used to deconstruct within-subject condition effects for both the brain and behavior metrics individually. All post hoc comparisons were corrected using Tukey adjustment. Outliers were identified by studentized residuals exceeding ±3. If removing the outlier observation did not change the statistical conclusions, then the original, full model was reported in the results (as in all cases but one). Non-normality of the dependent measures was evaluated using the Shaprio-Wilk test. A Box-Cox transformation procedure was performed on non-normal variables; specifically, the first convenient power parameter within the lambda confidence interval was used to select a transformation method (cf. optimal power parameter; specified by the “convenient” t-option). However, data transformations did not alter results for any non-normally distributed data, so the non-transformed results are reported below. Effect sizes are also included for the F-statistics and post hoc t-statistics using partial eta-squared (*η^2^p*) and Hedge’s *g* (repeated measures), respectively (Uanhoro, n.d.).

## Results

### ABR condition effects

#### Masking and speech in noise effects: single- vs. dual-talker

To evaluate speech-in-noise (SiN) effects on the ABR wave V, we compared Cheech responses for the male voice mono-talker and the dual-talker conditions using mixed effects regression analysis (i.e., main effect variable dummy coded for either mono- or dual-talker, where dual-talker was combined across male & female responses). Overall, the dual-talker condition was associated with both reduced wave V amplitudes (*F*_1,99_=25.84, *p*<0.0001, *η^2^p*=0.2070) and delayed latencies (*F*_1,99_=10.82, *p*=0.0014, *η^2^p*=0.0985) compared to the mono-talker condition. A one-way, repeated-measures ANOVA was then used to elucidate underlying relationships between all conditions individually (i.e., mono-talker male target, dual-talker male target, dual-talker male masker, dual-talker female target, and dual-talker female masker). Tukey-adjusted post hoc comparisons revealed larger amplitudes for the mono-talker (male) condition compared to all dual-talker conditions individually (*t*’s>3, *p*’s<0.05). Additionally, faster latencies were observed for the mono-talker condition compared to the dual-talker female target (Tukey-adjusted *t*_96_=-3.0020, *p*=0.0276, *g*=-0.4751) and dual-talker male masker (Tukey-adjusted *t*_96_=-3.6140, *p*=0.0043, *g*=-0.5790).

As mentioned in the methods, the study design included only a male speaker for the mono-talker condition (cf. female mono-talker) due to time constraints and excessive listening fatigue. Characteristic differences between the voices may potentially confound the results. A secondary analysis was therefore conducted to isolate the most similar, salient masking contrast: male mono-talker versus male dual-talker (target only). Consistent with the full model results, the male dual talker was associated with reduced wave V amplitudes (*F*_1,24_ =27.31, *p*<0.0001, *η^2^p*=0.5323) and delayed wave V latencies (*F*_1,24_=7.60, *p*=0.0110, *η^2^p*=0.2405) compared to the male mono-talker condition. Collectively, these results suggest poorer brainstem encoding for stimuli in the presence of a competing masker (i.e., smaller amplitudes and delayed latencies; Figure 3).

**Figure 3:**
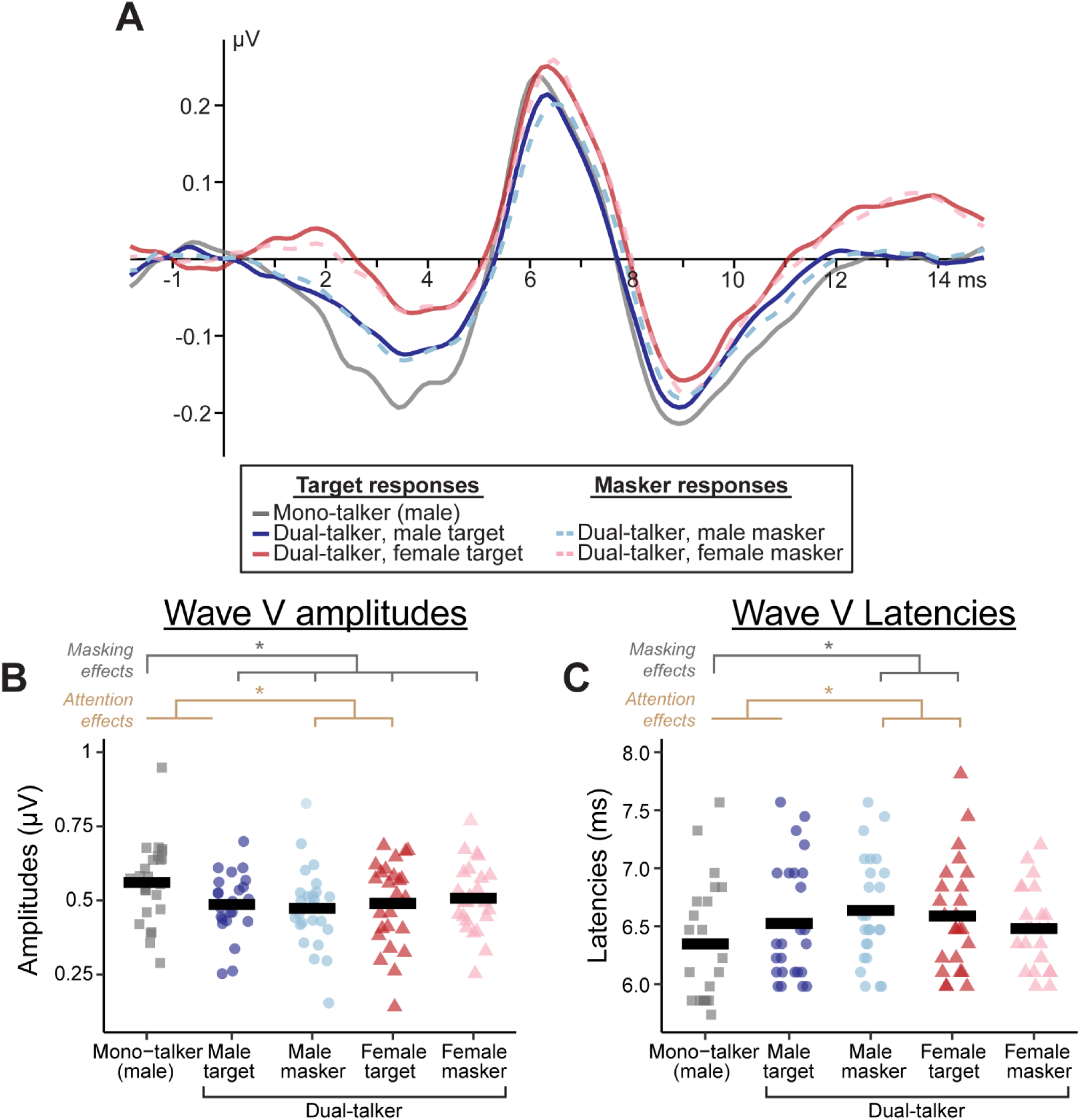
(A) ABR waveforms evoked by the Cheech chirps across target and masker conditions show wave V differences with the presence of another talker (speech-in-noise effects) as well as directed selective attention (target vs. masker). (B) Wave V amplitudes were larger for the mono-talker condition compared to all dual-talker conditions. The male target was also (combined mono- & dual-talker) was larger in amplitude compared to the male masker and female target. (C) Wave V latencies were faster for the mono-talker condition compared to the dual-talker male masker and female target conditions. Additionally, faster latencies were observed for the combined male target compared to the male masker and the female target voice. Figure icons depict individual subject observations while the bold black line indicates the mean. * = p<0.05.

#### Attention effects

Attention effects within the brainstem were assessed as the difference between target and masker responses across the listening blocks. Considering the effects of target versus masker alone, no differences were observed in either wave V amplitudes (*F*_1,99_=2.96, *p*=0.0885, *η^2^p*=0.0290) or latencies (*F*_1,99_=1.75, *p*=0.1886, *η^2^p*=0.0174). However, as suggested by the grand average waveforms (Figure 3A), the attention effect could be modulated by the talker’s voice. We therefore tested for the interaction of the talker sex and directed attention (target versus masker) as well as respective main effects on wave V measures. The interaction between selective attention and voice sex was significant for both wave V amplitudes (*F*_1,97_=7.19, *p*=0.0086, *η^2^p*=0.0690) and wave V latencies (*F*_1,97_=8.17, *p*=0.0052, *η^2^p*=0.0777). Tukey-adjusted simple effect comparisons suggested larger amplitudes for the male target (mono- and dual-talker combined) compared to the male masker (*t*_97_=3.03, *p*=0.0032, *g*=0.3712) and female target (*t*_97_=2.03, *p*=0.0447, *g*=0.2369). Additionally, faster latencies were observed for the male target compared to the male masker (*t*_97_=-2.84, *p*=0.0055, *g*=-0.3981) and the female target voice (*t*_97_=-2.15, *p*=0.0341, *g*=-0.2974) in post hoc contrasts.

Although the mono-talker male could be considered a target condition itself, even in the absence of a masker talker, it is possible that including this condition in the models introduces a confound in the statistical models. Like the *Masking effects* results above, we thus performed a secondary analysis on a subset of the data that only included the male and female dual-talker conditions (i.e., both target and masker). With the male mono-talker removed, for wave V amplitudes neither target versus masker nor male versus female voice main effects were significant (*F*_1,72_=0.03 & 2.02, *p*=0.8585 & 0.1593, *η^2^p*=0.0004 & 0.0273, respectively) nor was their interaction (*F*_1,72_=1.36, *p*=0.2480, *η^2^p*=0.0185). However, the interaction was borderline significant after removal of one outlier observation (i.e., from the male masker condition; *F*_1,71_=3.65, *p*=0.0601, *η^2^p*=0.0489), and the main effect of voice sex suggested slightly smaller amplitudes for the dual-talker male compared to the female voice (*F*_1,71_=4.87, *p*=0.0306 *η^2^p*=0.0642). Meanwhile, the interaction was borderline for wave V latencies (*F*_1,72_=3.66, *p*=0.0598, *η^2^p*=0.0484). These results suggest the attention effects described above for the full models are at least partially driven by larger amplitudes and faster latencies for the mono-talker male condition compared to the dual-talker conditions, but a general trend for attention effects in the auditory brainstem is still evident.

### Behavioral performance

One-way repeated measures ANOVAs were used to evaluate differences in behavioral performance metrics across conditions (i.e., mono-talker male, dual-talker male, & dual-talker female). Selective attention performance was measured behaviorally as identification accuracy and corresponding reaction times for color words embedded in the target story (Figure 4). Color word identification accuracy significantly differed across all three experimental conditions (*F*_2,48_=18.15, *p*<0.0001, *η^2^p*=0.4306; Figure 4A). Specifically, both the mono-talker male and dual-talker male conditions were associated with higher identification accuracy than the dual-talker female (*t*_48_=4.9768 & *t*_48_=5.4303, *g*=0.7903 & *g*=0.8563, respectively; both Tukey-adjusted p<0.0001). The two male talker conditions were not significantly different (*t*_48_=-0.4534, *p*=0.8931, *g*=-0.1030), suggesting a slight behavioral identification accuracy decrement specific to the female-voiced story, not necessarily with the general presence of a competing talker. Meanwhile, target color word reaction times also differed across conditions (*F*_2,48_=10.02, *p*=0.0002, *η^2^p*=0.2945). Specifically, faster reaction times were observed for the mono-talker block compared to the dual-talker female (*t*_48_=-4.4761, *p*=0.0001, *g*=-0.6725; Figure 4B). Although the other Tukey-adjusted post hoc comparisons were not significant, reactions times trended towards faster reaction times for the mono-talker male compared to the dual-talker male (*t*_48_=-2.2442, *p*=0.0739, *g*=-0.3349) and for the dual-talker male compared to the dual-talker female condition (*t*_48_=-2.2318, *p*=0.0760, *g*=-0.3177).

**Figure 4:**
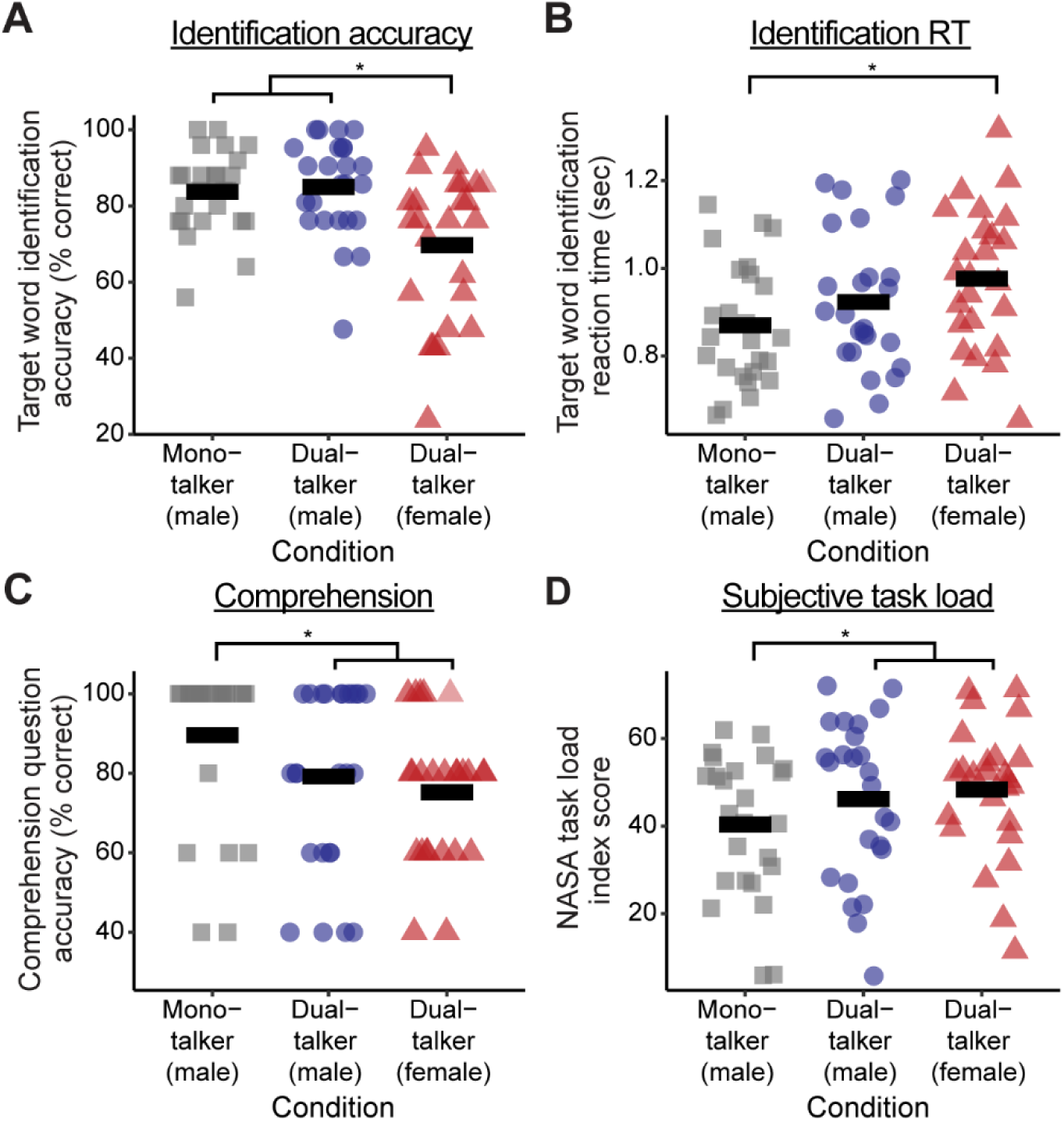
Behavioral results for color word identification accuracy (A), identification reaction times (B), narrative comprehension (C), and ratings of subjective task load or effort (D). Figure icons depict individual subject observations while the bold black line indicates the mean. * = p<0.05.

Following the listening task, participants were asked to complete several comprehension questions to probe their understanding of the short story segments. Response accuracy for comprehension questions was higher for the mono-talker condition compared to either dual-talker condition (overall *F*_2,48_=5.97, *p*=0.0048, *η^2^p*=0.1991; Tukey-adjusted mono-talker vs. dual-talker male *t*_48_=2.4175, *p*=0.0501, *g*=0.4681 & mono-talker vs. dual-talker female *t*_48_=3.3473, *p*=0.0045, *g*=0.7323; Figure 4C). No difference was observed between the male- and female-voiced dual-talker conditions (*t*_48_=0.9298, *p*=0.6243, *g*=0.1897). Participants then completed the NASA Task Load Index questionnaire (Hart and Staveland, 1988) as a measure of their subjective task load or effort. The mono-talker story was associated with lower scores (i.e., less perceived task load) compared to either dual-talker condition (dual-talker male: *t*_48_=-2.8322, *p*=0.0182, *g*=-0.3204; dual-talker female: *t*_48_=-3.8978, *p*=0.0009, *g*=-0.4899), but the dual-talker blocks did not significantly differ from each other (*t*_48_=-1.0636, *p*=0.5410, *g*=-0.1227; Figure 4D). Collectively, the results suggest that the dual-talker scenario was associated with both greater perceived effort (task load) and impaired comprehension abilities.

Relationships between behavioral measures were also evaluated using linear mixed-effects regression models. All four behavioral outcomes were associated with each other (all *p*’s < 0.05). For further details and individual statistical analyses, see Extended Data Figure 4-1. As might be expected, better behavioral performance was characterized by higher identification accuracy, faster identification reaction times, higher comprehension question accuracy, and lower effort ratings.

### Brainstem-behavior relationships

We first analyzed mixed effects models with main effects and interactions of directed attention (i.e., target versus masker), talker sex, and wave V measures to predict behavioral outcomes (i.e., separate models for latencies and amplitudes). No 3- or 2-way interactions were observed between either wave V measure and directed attention or talker sex variables (all p’s >0.05). The only 2-way interactions between talker sex and directed attention observed in the full model simply replicated findings previously reported in the Behavioral Performance section.

The models were then re-analyzed to assess only main effects of ABR wave V amplitudes and latencies separately on behavioral outcomes (Figure 5 & Table 1). These exploratory analyses broadly investigate whether wave V across all the conditions relates to behavioral outcomes rather than being tied to specific condition effects. In these simplified analyses, faster wave V latencies were associated with better target word identification accuracy (*F*_1,99_=8.59, *p*=0.0042, *η^2^p*=0.0798), better comprehension question accuracy (*F*_1,99_=4.73, *p*=0.0320, *η^2^p*=0.0456), and lower subjective task load (*F*_1,99_=19.28, *p*<0.0001, *η^2^p*=0.1630). Latencies alone did not correlate with target word identification reaction times (*F*_1,99_=1.06, *p*=0.3050, *η^2^p*=0.0106). Further, larger wave V amplitudes were associated with better target word identification accuracy (*F*_1,99_=9.52, *p*=0.0026, *η^2^p*=0.0877) and reaction times (*F*_1,99_=8.82, *p*=0.0037, *η^2^p*=0.0818) but not comprehension question accuracy nor subjective task load (*F*_1,99_=1.41, *p*=0.2377, *η^2^p*=0.0140 & *F*_1,99_=0.66, *p*=0.4172, *η^2^p*=0.0066, respectively). Collectively, these results indicate a strong relationship between wave V and measures of selective attention (i.e., target word identification). Meanwhile, neural encoding speed (i.e., wave V latencies) but not encoding strength (i.e., amplitudes) related to both narrative comprehension and perceived effort.

**Figure 5:**
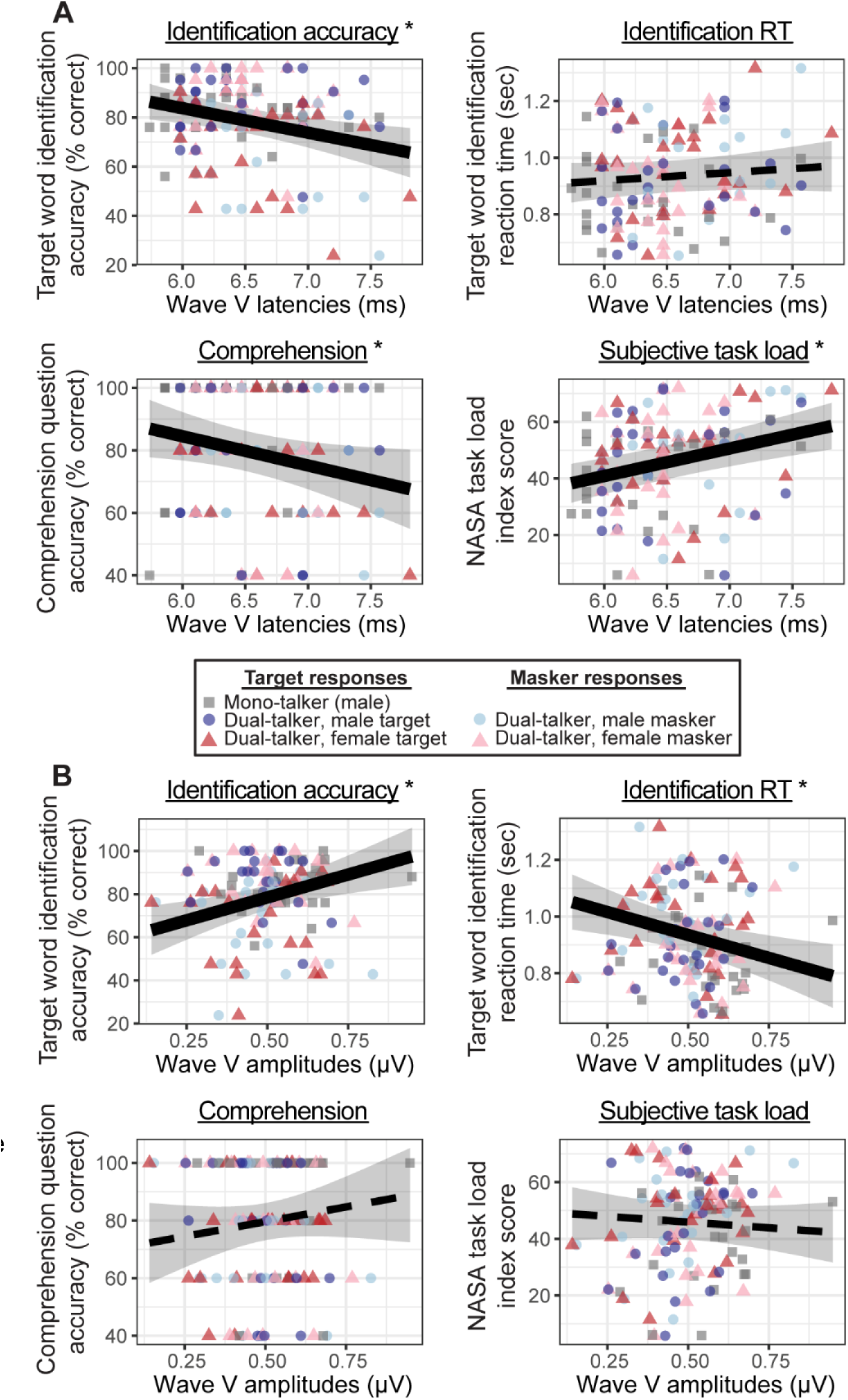
Associations between auditory brainstem response (ABR) wave V (A) amplitudes and (B) latencies and each of the four behavioral metrics. The figures show individual observations across all subjects and conditions with the regression line depicting the brain-behavior relationship. Bolded regression lines and an asterisk in the plot title indicate significant fixed effect results (p<0.05), whereas dotted lines are not significant (p>0.05). See Table 1 for regression results. Shaded region = ± 95% CI.

**Table 1:** Brainstem-behavior regression model results for wave V main effects.

| Wave V predictor | Outcome | $\beta$ estimate | $F_{1,99}$ statistic | 95% Confidence interval | P-value |
| --- | --- | --- | --- | --- | --- |
| Latencies | Target word identification accuracy (%) | -9.9569 | 8.59 | (-16.6981, -3.2158) | **0.0042\*** |
|  | Target word identification reaction times (sec) | 0.0283 | 1.06 | (-0.0261, 0.0828) | 0.3050 |
|  | Comprehension question accuracy (%) | -9.3495 | 4.73 | (-17.8770, -0.8221) | **0.0320\*** |
|  | Subjective task load scores | 9.7502 | 19.28 | (5.3446, 14.1558) | **<0.0001\*** |
| Amplitudes | Target word identification accuracy (%) | 42.4089 | 9.52 | (15.1361, 69.6816) | **0.0026\*** |
|  | Target word identification reaction times (sec) | -0.3235 | 8.82 | (-0.5397, -0.1074) | **0.0037\*** |
|  | Comprehension question accuracy (%) | 20.4951 | 1.41 | (-13.7387, 54.7289) | 0.2377 |
|  | Subjective task load scores | -8.0859 | 0.66 | (-27.7779, 11.6062) | 0.4172 |

Faster wave V peak latencies and larger amplitudes were associated with enhanced identification of the target words embedded within the stories. Faster latencies additionally related to better comprehension question accuracy and lower subjective listening effort. \*=p<0.05

## Discussion

We evaluated the effects of speech-in-noise (SiN) and selective attention on low-level brainstem encoding. Our novel experimental paradigm permitted simultaneous measurement of both ABRs and behavior in both single- and dual-talker conditions. ABRs recorded in the presence of a competing talker were associated with smaller amplitudes and delayed latencies compared to the mono-talker condition. Interestingly, the results also suggested a role of directed attention on ABR wave V latencies. These effects on directed attention were partially driven by talker’s sex and mainly occurred in the context of the mono-talker male voice as a target condition. Faster wave V latencies were associated with better accuracy of color word identification as well as higher comprehension question accuracy and lower scores of subjective listening effort. Meanwhile, more robust wave V amplitudes corresponded to better accuracy and faster reaction times for target word identification during the selective attention listening task. Collectively, the results suggest that faster, more robust neural encoding within the auditory brainstem may contribute to successful auditory selective attention and SiN listening processes.

Selective auditory attention within the brainstem was characterized by wave V response differences when participants were instructed to attend to or ignore short story narratives. However, this effect was restricted to analyses where the mono-talker male condition was included as a target condition, thus including any differences between mono-versus dual-talker masker conditions. Yet, these results align with other reports of attention-driven influences on subcortical processing. For example, Forte et al. (2017) and Saiz-Alía & Reichenbach (2020) noted larger computationally-derived auditory brainstem amplitudes for attended over unattended speech. Other studies have shown modulation of transient-evoked ABRs while attending to the auditory modality versus another (typically visual) domain (Lukas, 1980; Bauer and Bayles, 1990; Ikeda and Campbell, 2021, 2024; Kumar et al., 2023) or with increasing cognitive interference (Sörqvist et al., 2012; Brännström et al., 2020). Our results are also consistent with attentional modulation of sustained subcortical frequency-following responses (Hoormann et al., 2000; e.g., Galbraith et al., 2003; Hairston et al., 2013; Lehmann and Schönwiesner, 2014; cf. Varghese et al., 2015). It is likely that corticofugal pathways— particularly those projecting to the inferior colliculus, a key neural generator of wave V (Starr and Hamilton, 1976; Hashimoto et al., 1981; Huffman and Henson, 1990)—contribute to directed attention effects on the ABR. Yet, the ABR has historically been viewed as largely immune from attention effects (e.g., Amadeo and Shagass, 1973; Picton and Hillyard, 1974; Connolly et al., 1989; Gregory et al., 1989). These discrepancies across studies likely result from differences in experimental paradigms. It is possible that attention-related effects on subcortical processes are most effectively revealed when another attention-demanding task is presented concurrently with the auditory stimuli (Lukas, 1980; e.g., Brännström et al., 2020; Kumar et al., 2023) or when the stimuli are more ecologically valid and relevant for real-world listening, as in the case of continuous speech (Ding and Simon, 2012; Zion Golumbic et al., 2013; Forte et al., 2017; Backer et al., 2019; Etard et al., 2019; Teoh and Lalor, 2019; Saiz-Alía and Reichenbach, 2020; Teoh et al., 2022; Carta et al., 2024; Shehabi et al., 2025). Thus, Cheech demonstrates potential for reliable detection of top-down, selective attention effects on subcortical processes.

In our exploratory analyses, faster and/or larger Cheech-evoked wave V responses were associated with more successful speech listening performance, including behavioral measures of selective attention (i.e., identification accuracy, RTs) as well as comprehension questions and subjective reports of listening effort. That is, more robust encoding in lower-level brainstem generators was related to higher-level speech recognition abilities, with and without the presence of a competing talker. Subcortical responses have been associated with other speech-listening behaviors including SiN perception (Papakonstantinou et al., 2011; Bramhall et al., 2015; Saiz-Alía and Reichenbach, 2020), self-reports of speech-listening abilities (Anderson et al., 2013), early categorization of speech phonemes (Carter and Bidelman, 2023; Rizzi and Bidelman, 2023), or even measures of auditory processing disorders (Omidvar et al., 2023). Collectively, our results provide further support for the important role of the brainstem in speech perception, both in quiet and the presence of a masking talker, as well as higher-level narrative comprehension and subjective perception of task-related listening effort.

As expected, the presence of a competing talker was associated with weaker brainstem encoding (i.e., wave V reduced amplitudes, delayed latencies), worse comprehension accuracy, and higher ratings of subjective task load. SiN perception (the “cocktail party problem”) is an important ability for listening in real-world environments, and some speech-hearing difficulties are only exposed when stimuli are presented in background noise. It remains possible that the variability of SiN perceptual skills, even among normal hearing listeners, could be traced to low-level neural encoding differences (e.g., Bharadwaj et al., 2015; Bramhall et al., 2015, 2017; Jain et al., 2019) or even top-down corticofugal modulations (Price and Bidelman, 2021). Varied contributions (or degeneration/selective loss) of auditory nerve fibers during speech encoding, particularly those with low-spontaneous discharge rates, could also help explain individual SiN differences (Mehraei et al., 2016, 2017). While the current study cannot differentiate between these theories—since both top-down and/or bottom-up mechanisms could alter ABR wave V and its relationship with speech perception—our results provide evidence of subcortical involvement for continuous SiN and suggest a possible neural factor for individual perception differences.

Contrasting with our findings, previous studies that compared male and female talkers reported larger wave V response amplitudes for male voices compared to female voices (Forte et al., 2017; Saiz-Alía and Reichenbach, 2020; Polonenko and Maddox, 2021). Yet, our selective attention-talker sex interaction for wave V amplitudes did align with previous reports of larger male-attended amplitudes compared to unattended speech, though a similar trend was noted for the female voice in Forte et al. (2017). One explanation could be due to differences in experimental stimulus design. Our Cheech and/or female-voice modulation processes could have affected perceptual clarity and thus impacted brain and/or behavioral responses even though average RMS values were equated across the stories. Our female-sounding voice was modified from the original male voice (same talker for all stories) to maintain similar intonation, phrasing, and other expressive characteristics across the stories; only the speaker’s F0 and vocal tract length parameters were resynthesized using MATLAB STRAIGHT to create a female-sounding voice (Kawahara and Morise, 2011). The voice modulation may also have impacted the relative chirp-to-speech energy leading to differences across the male and female talkers in this study. Additionally, higher rates of stimulation are associated with reduced ABRs (e.g., Don et al., 1977; Weber and Fujikawa, 1977; Fowler and Noffsinger, 1983). Higher F0s and thus faster glottal pulse rates could explain subcortical encoding differences from speech stimuli spoken by different talkers (Polonenko and Maddox, 2021). While our stimuli had approximately 10% more chirps for female-voiced stories compared to the male-voiced stories (i.e., faster glottal pluses creates more chirp-embedding opportunities), we set a minimum inter-stimulus interval of 18.2 ms between chirps to reduce overlapping activity and detrimental effects of high stimulation rates on neural encoding processes. We also equated the number of chirp trials across conditions in our analyses to avoid any systematic confounds produced by talker sex-based SNR differences due to imbalanced chirp events. Finally, chirps are also designed to activate the entire cochlea in a near-synchronous fashion which produces more robust ABRs in general (Elberling et al., 2007, 2010) which could explain neurophysiologic differences between our study and other computationally-derived methods more generally.

### Limitations

Our male and female talkers were spatially separated using head-related transfer functions (HRTFs), and the talker locations were dynamically swapped with each other on average every 6 seconds to track spatially-driven selective attention similar to Teoh & Lalor (2019). Pilot testing confirmed that the combined effects of pitch modulation (to produce a female-sounding voice) and spatial separation led to better listening performance, consistent with previous literature demonstrating talker-sex, spatially-separated release of masking effects in multi-talker environments (Oh et al., 2021, 2022). However, gaze direction (as when participants were asked to follow the target cue across the two screens) could potentially modulate neural or myogenic activity that could contaminate responses (e.g., O’Beirne and Patuzzi, 1999; Patuzzi and O’Beirne, 1999; Bidelman et al., 2024). In our experiment, only male-spoken stories were included for the mono-talker condition. Female mono-talker stories were omitted from the final experiment after pilot testing to reduce the total testing time and potential fatigue. Participants therefore receive more exposure to the male voice during the speech listening task. Additionally, attention effects on the full dataset (imbalanced design with no female mono-talker) did not wholly replicate for data subsets that were more balanced across conditions (i.e., dual-talker only), though the statistics suggested a similar albeit potentially underpowered underlying pattern in the subset data. Thus, we cannot rule out certain idiosyncratic effects of our experimental paradigm on the observed brainstem-behavior relationships. Different experimental designs may lend better to other types of analyses, such as mediation analyses, that may shed more light on whether brainstem responses mediate the relationship between speech-listening conditions and behavioral outcomes. Future studies, both involving Cheech and other (ideally continuous) speech stimuli, may help elucidate the underlying neural encoding mechanisms involved in processing individual talker characteristics and its impact on speech perception abilities.

## Conclusion

Collectively, our results provide evidence for the role of brainstem encoding process in individual speech perception abilities including SiN recognition, selective auditory attention, and comprehension. These findings underscore the importance of lower-level auditory structures for speech processing, even in audiometrically normal listeners. They furthermore have implications for an improved understanding of other populations (e.g., older adults, children, individuals with hearing loss or assistive hearing devices, etc.) and of certain speech processing disorders. Our study highlights the advantage of a stimulus like chirped-speech (Cheech) and more ecologically valid paradigms to uncover novel and impactful relationships between lower-level neural processing and higher-level listening skills in real-world auditory environments.

## Extended Data

**Extended Data Table 1-1.** Short story stimulus details.

| **Story title** | **Author** | **Collection** | **Experimental condition** | **Male or female voice** |
| --- | --- | --- | --- | --- |
| The Master Cat, or Puss in Boots | Charles Perrault, translated by J. E. Mansion and Robert Samber | The Fairy Tales of Charles Perrault | Mono-talker | Male |
| Lily and the Lion | Jacob Grimm & Wilhelm Grimm | Grimms' Fairy Tales | Dual-talker | Male |
| The Four Clever Brothers | Jacob Grimm & Wilhelm Grimm | Grimms' Fairy Tales | Dual-talker | Female |
| The Wanderers | Edith Howes | Maoriland Fairy Tales | Practice only (mono-talker) | Male |

**Extended Data Table 1-2.** Cheech synthesis parameters.

| **<u>Cheech Parameter</u>** | **<u>Value</u>** |
| --- | --- |
| Fs (sample rate; Hz) | 48000 |
| Speech filter bands (Hz) | 0-250; 500-1000; 2000-4000; 11000+ |
| Chirp filter bands (Hz) | 250-500; 1000-2000; 4000-11000 |
| Speech filter order | 10 |
| Chirp filter order | 4 |
| Voicing (speech envelope) filter, to identify voicing by power (Hz) | 20-1000 |
| Voicing threshold, for identifying sufficient glottal pulse energy (normalized amplitude) | 0.03 |
| Minimum voiced period to chirp (sec) | 0.05 |
| Minimum chirp ISI (sec) | 0.0182 |
| Minimum gap after first chirp (sec) | 0.048 |
| Maximum voicing duration, above which gaps will be inserted (intra-voicing gap) to prevent very long chirp trains especially during loud speech modulations (sec) | 0.400 |
| Intra-voicing gap (sec) | 0.05 |
| Normalized chirp amplitude | 0.10 |
| Female voice characteristics<br>Resynthesized voice mean F0 pitch (Hz) | 180<br>4000 |
| Maximum frequency modulation (Hz) | 1.04 |
| Vocal tract length warping factor |  |
| Resynthesized voice characteristics |  |
| Score, to reduce thumping in the resynthesized audio | 0.6 |
| Margin, to reduce buzzing in the resynthesized audio | 0.6 |

**Extended Data Figure 4-1:**
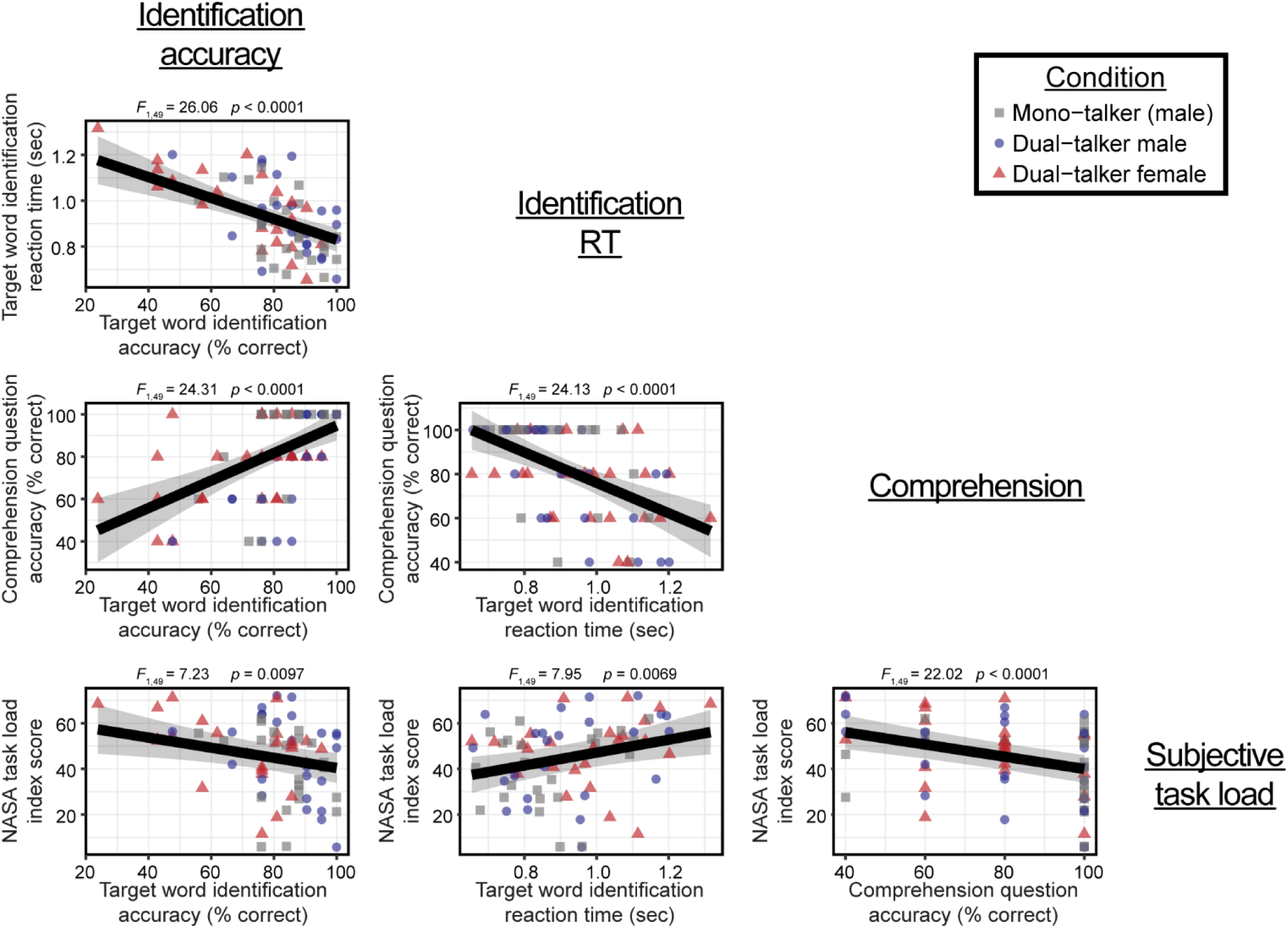
Associations between the four behavioral metrics. Each figure plots the individual observations across the three target conditions (mono-talker male, dual-talker male, and dual-talker female) and the regression line depicts the relationship between the behavioral variables. Specifically, better behavioral performance was characterized by higher identification accuracy, shorter identification reaction times (RT), higher comprehension accuracy, and lower subjective task load or effort (NASA task load index scores). Mixed effects regression results for the main fixed effect are shown at the top of each figure plot. Shaded region = ± 95% confidence interval.

## Acknowledgements

The authors would like to thank members of the University of California, Davis Health audiology and clinical research team for performing initial hearing evaluations of our participants: Dr. Robert Ivory, AuD; Dr. Mackenzie Quinn, AuD; Dr. Rachel Krager, AuD; Dr. Steven Zurawski, AuD; Dr. Austin Childers, AuD; Dr. Kimberly Smith, AuD; Angela Beliveau, and Randev Sandhu. We are also grateful for our team of undergraduate lab volunteers—particularly Jillian McKie and Cathleen Chan—who assisted in countless hours of participant recruitment, data collection, and data pre-processing. Thank you to Tiana Smith and Elyse Ehlert for their efforts in participant recruitment.

AI was used in a limited capacity for this work, primarily assisting with minor text editing and basic phrasing. All significant drafting, analysis, and editing were carried out by the authors.

## Dedications

This research is dedicated in loving memory of our research team member, fellow lab mate and friend, Karim Abou Najm.

